# A Molecular Mechanism for Probabilistic Bet-hedging and its Role in Viral Latency

**DOI:** 10.1101/2020.05.14.096560

**Authors:** Sonali Chaturvedi, Jonathan Klein, Noam Vardi, Cynthia Bolovan-Fritts, Marie Wolf, Kelvin Du, Luwanika Mlera, Meredith Calvert, Nathaniel J. Moorman, Felicia Goodrum, Bo Huang, Leor S. Weinberger

## Abstract

Probabilistic bet hedging, a strategy to maximize fitness in unpredictable environments by matching phenotypic variability to environmental variability, is theorized to account for the evolution of various fate-specification decisions, including viral latency. However, the molecular mechanisms underlying bet-hedging remain unclear. Here, we report that large variability in protein abundance within individual herpesvirus virion particles enables probabilistic bet hedging between viral replication and latency. Super-resolution imaging of individual virions of the human herpesvirus cytomegalovirus (CMV) showed that virion-to-virion levels of pp71 tegument protein—the major viral transactivator protein—exhibit extreme variability. This super-Poissonian tegument variability promoted alternate replicative strategies: high virion pp71 levels enhance viral replicative fitness but, strikingly, impede silencing, whereas low virion pp71 levels reduce fitness but promote silencing. Overall, the results indicate that stochastic tegument packaging provides a mechanism enabling probabilistic bet hedging between viral replication and latency.

**SIGNIFICANCE:** Probabilistic bet hedging is a generalized diversification strategy to maximize fitness in unpredictable environments, and has been proposed as an evolutionary basis for herpesvirus latency. However, the molecular mechanisms enabling probabilistic bet hedging have remained elusive. Here, we find that the human herpesvirus cytomegalovirus—a major cause of birth defects and transplant failures—utilizes stochastic variability in the abundance of a protein packaged into individual viral particles to enable probabilistic bet hedging between alternate viral states.

## INTRODUCTION

Diverse biological systems share a common challenge to preserve reproductive fitness in unpredictable, changing environments. Faced with environmental variability, some organisms probabilistically generate a range of phenotypes to ‘hedge their bets’ (1-7), in much the same way that financial houses diversify their assets to minimize risk against economic crashes. First proposed over 50 years ago (1) for desert annuals— where reproductive success is subject to unpredictable weather patterns—bet-hedging theory noted that temporal variation in fitness could be minimized if husk thickness between seeds varied such that random chance biased the population’s germination potential. In this way, some seeds randomly enter dormancy irrespective of the environment and a long-lived, desiccation-resistant subpopulation is always formed to avoid extinction during unforeseen droughts, but at the necessary expense of lowering germinative fitness. In the decades since, bet hedging has been studied in persistence phenotypes in bacteria (2, 5, 7, 8), yeasts (9, 10), and viruses (11-13). In herpesviruses, bet-hedging theory has been proposed as a theoretical basis for the evolution of viral latency (11) and associated viral gene silencing. However, the molecular mechanisms that allow biological systems, such as viruses, to probabilistically generate the needed variability have remained unclear and an area of active study.

In the herpesviridae family, persistence within the host is mediated by establishment of a reversible latent state, wherein viral gene expression is largely silenced (14, 15). In CMV—one of nine human herpesviruses and a leading cause of birth defects and transplant failure—lytic replication occurs in a variety of cell types, while latency and requisite silencing are established in myeloid-progenitor cells (14, 16, 17). The silent state is thought to enable evasion of host immune responses and confer a selective advantage (11). To reactivate from a silenced state or initiate lytic replication, expression from CMV’s Major Immediate-Early Promoter (MIEP) is essential. During silencing, the MIEP is largely quiescent, but during lytic replication the MIEP drives expression of crucial viral genes including the 86-kDa Immediate-Early 2 (IE2) protein, a master regulator of lytic viral expression. Initiation of lytic replication requires that the MIEP be efficiently transactivated by proteins carried within the viral tegument (18), a proteinaceous region between the viral capsid and envelop. The pp150 (UL32) and pp71 (UL82) tegument proteins appear to be the chief tegument regulators of MIEP expression (19-21), with pp71 transactivating MIEP activity and pp150 repressing MIEP activity.

Building off previous quantitative analyses of tegument variability in other herpesviruses (22-24), we quantified CMV single-virion variability in pp71 and pp150 levels using super-resolution fluorescence microscopy, and then examined how variability in particle-associated protein abundance affects CMV replicative strategies. We find that virion-associated pp71 levels are significantly more variant than pp150 levels or than expected by Poisson statistics. When virions with increased pp71 levels were generated, the population exhibited enhanced infectiousness and replicative fitness. This enhanced fitness conferred by higher pp71 raised the question of why selective pressures had not forced the pp71 distribution to a higher mean and lower variance, indicating that a putative counterbalancing selection pressure may exist to maintain the broad pp71 distribution. Indeed, we found that low pp71 levels promoted MIEP silencing whereas high pp71 levels impeded silencing in undifferentiated cells. We propose a conceptual model for how heterogeneity in the levels of virion-packaged tegument transactivators may enable probabilistic bet hedging between replication and silencing in herpesviruses.

## RESULTS & DISCUSSION

### The HCMV major tegument transactivator protein pp71 exhibits super-Poissonian variability in virion-to-virion abundance

To measure the variation in pp71 and pp150 abundance in individual virion particles, we utilized a super-resolution imaging method (25) that enabled fluorescent imaging of pp71 or pp150 tegument proteins that were genetically tagged with yellow fluorescent protein (YFP) (**Fig. 1*A***). To obtain the viral samples for imaging, recombinant virus was packaged in culture and viral preparations were gradient purified via ultracentrifugation to isolate infectious particles and exclude dense bodies and empty particles. Electron microscopy verified the purity of single, intact virions (*SI Appendix*, **Fig. S1**). For image quantification, we used an established super-resolution segmentation algorithm (26) that ensures only singlet particles are analyzed, and scored and implemented segmentation based on the lipid envelope co-staining to ensure that only envelope-associated particles were analyzed (*SI Appendix*, **Fig. S2**). Only virus-sized (∼200nm diameter) singlet particles were included in the imaging analysis, and a minimum of 2000 intact, singlet particles were quantified for each fusion virus. To control for contributions from instrument and photon shot noise, a ‘molecular ruler’ (27), containing precisely 900 florescent-fused HSV-1 VP26 proteins per viral particle (24), was imaged in parallel and used for comparison.

**Figure 1:**
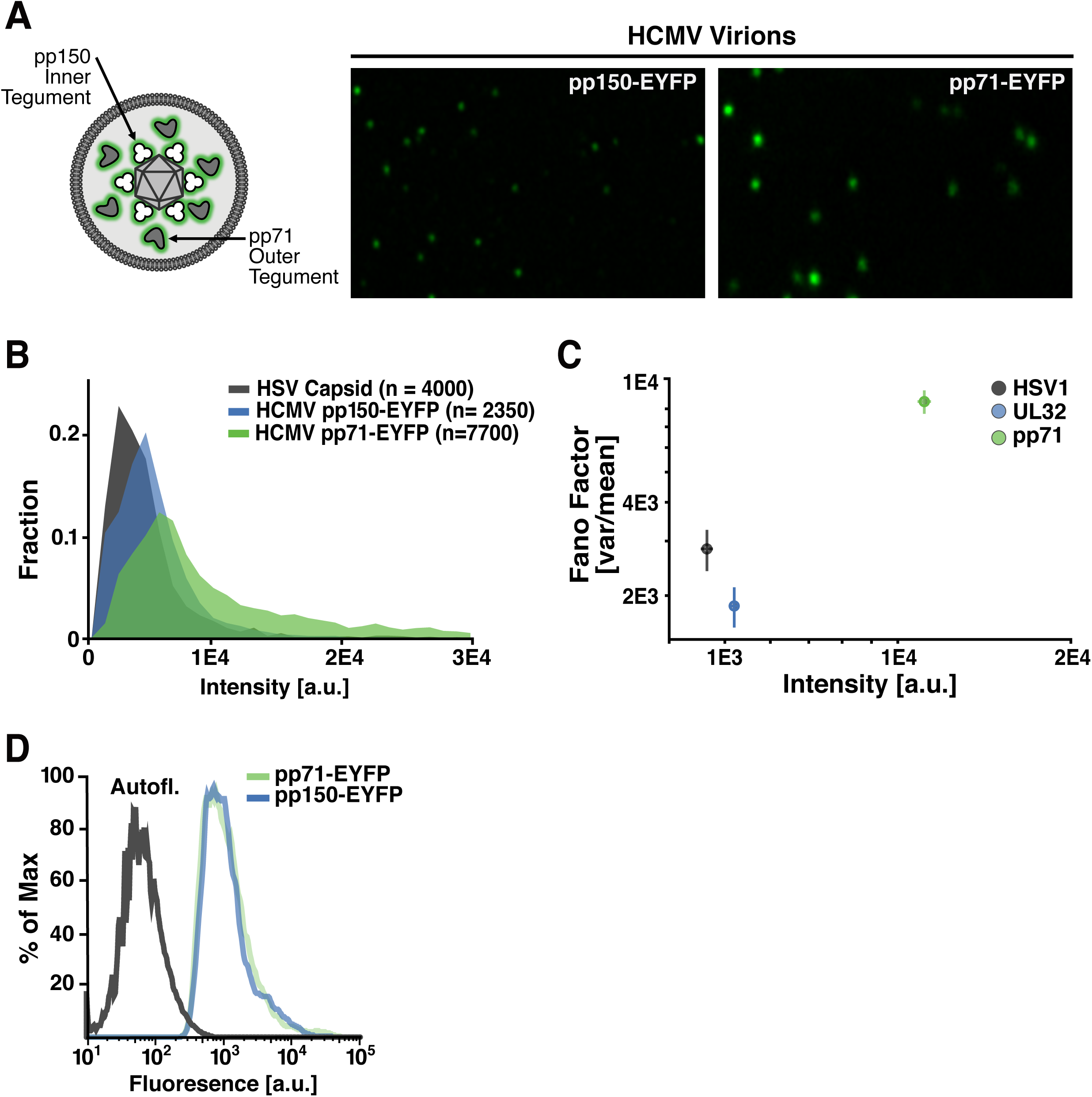
Substantial virion-to-virion heterogeneity in the level of the HCMV major tegument transactivator protein pp71. (*A*) (Left) Schematic of HCMV virion showing relative locations of pp150 (UL32), which is capsid-associated, in the inner tegument, and pp71 (UL82), which is capsid-unassociated, in the outer tegument. (Right) Representative super-resolution fluorescence micrographs of purified, infectious, recombinant HCMV virion particles where tegument proteins are genetically fused to EYFP: pp150-EYFP (left), pp71-EYFP (right). (*B*) Quantification of pp71 and pp150 abundance in individual virion particles relative to the HSV-1 VP26-GFP ‘molecular ruler’ (900 copies per virion). (*C*) Normalized variance (Fano factor; σ^2^/μ) versus mean abundance (<intensity>) in pp71, pp150 and HSV-1 capsid in the virion populations. Error bars were estimated by bootstrapping the data with n=1000 particles per sample, 150 times. (*D*) Intracellular levels of tegument factors EYFP-pp71 and pp150-EYFP in infected human fibroblasts quantified by flow cytometry (left panel).

Quantitative image analysis showed that the per virion levels of the pp150 tegument protein, which is capsid associated, fell within a relatively confined range, exhibiting 2−3 fold particle-to-particle variation in intensity. (**Fig. 1*B***). This variation in pp150 was similar to the virion-to-virion variation in HSV-1 VP26 capsid levels (**Fig. 1*B***), which are considered invariant (24). However, in striking contrast, pp71 tegument levels imaged under identical conditions and parameters varied >10-fold between the dimmest and brightest virion particles (**Fig. 1*B***), with the normalized virion-to-virion variance (σ^2^/μ, a standardized measure of the histogram width referred to as the ‘Fano factor’) being 5-fold greater for pp71 than pp150 (**Fig. 1*C***). Importantly, an orthogonal confocal imaging method showed close agreement with the super-resolution imaging measurements (*SI Appendix*, Fig. S2B). Moreover, these image-based quantifications of pp71 and pp150 are within the range of previously reported ratios from mass-spectrometry analyses (28).

To determine whether the large pp71 particle-to-particle variability resulted from a subpopulation of virus-producing cells that are ‘outliers’ and express higher levels of pp71 but not pp150—or other extrinsic variability in the virus-producing cells—we analyzed pp71 and pp150 expression levels in virus-producing cells using flow cytometry. In contrast to virion-particle heterogeneity, pp71 and pp150 expression levels in virus-producing cells exhibited only minimal cell-to-cell variability that could not account for the relative amplification of pp71 heterogeneity in virion particles (**Fig. 1*D***). While pp71 expression levels did not vary between cells, the distribution of pp71 puncta *within* virus-producing cells—assayed by super-resolution imaging—showed high variability. Specifically, cytoplasmic pp71 foci that were virion sized (i.e., smaller than 300nm) showed a long-tailed distribution in intensity with a large normalized variance (σ^2^/μ = 16782) (*SI Appendix* **Fig. S2C**). While the distribution of pp150 puncta *within* virus-producing cells also exhibited similar variability, the capsid-associated nature of pp150 appears to effectively filter this heterogeneity since overexpression of pp150 in cells did not increase pp150 levels in virions (*SI Appendix*, **Fig. S2C**), whereas pp71 overexpression in cells did increase pp71 in virions (18). These results argue that intrinsic heterogeneity in the spatial distribution of intracellular pp71, rather than extrinsic heterogeneity arising from expression-level differences between infected cells, contributes to virion-to-virion variability in pp71 levels.

### High pp71 offers an evolutionary advantage by increasing infectivity

To test if this large inter-virion pp71 variability carried a functional consequence, we generated CMV virion particles that packaged increased levels of pp71 protein (pp71^HI^) using a cell line expressing pp71, as previously described (29). Importantly, pp71 virion levels can be modulated without affecting packaging of other tegument factors (30). Super-resolution image analysis verified that singlet pp71^HI^ virions contained substantially more pp71, on average ∼2-fold higher than the pp71 levels in pp71^WT^ virus produced from standard fibroblasts (**Fig. 2*A***). In contrast, when virus was packaged on cells over-expressing pp150, no measurable increase in virion pp150 levels could be detected (*SI Appendix*, **Fig. S2C**), presumably because pp150 is capsid-associated and its levels fixed by the invariable number of capsid proteins.

**Figure 2:**
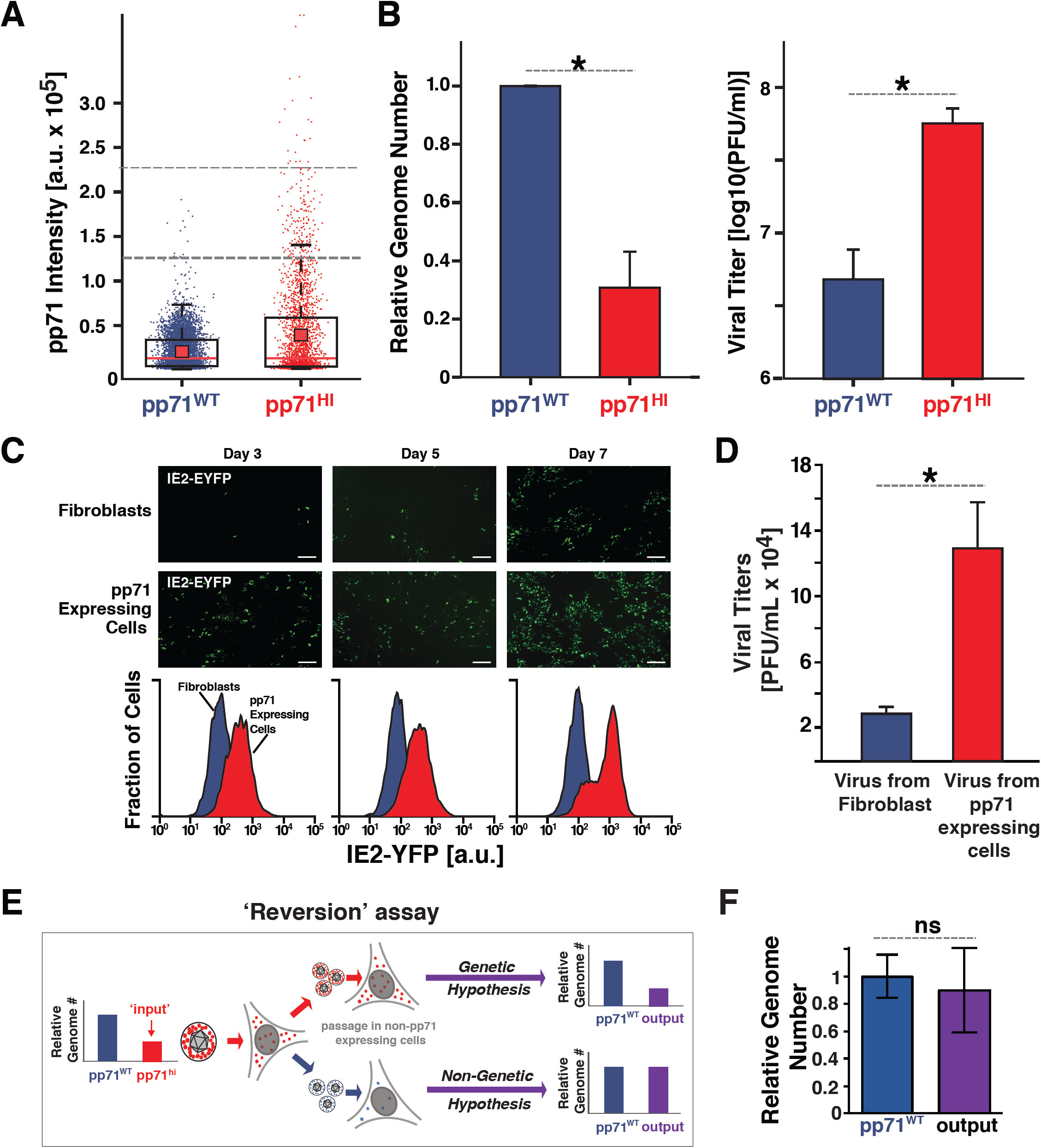
The ‘advantage’ of high pp71: High virion pp71 levels enhance HCMV replicative fitness. (*A*) Box-whisker plot of pp71 levels (by EYFP intensity) in purified virion particles quantified by super-resolution microscopy: pp71^WT^ EYFP virion particles (purple, n=5519); pp71^HI^ EYFP virion particles (red, n=2463). Analysis was restricted to particles 100−300nm in diameter. The red square depicts the mean μ, red line the median, the box encloses values between the first and third quantiles (25− 75%), and the whiskers (error bars) show the min and max intensities. The horizontal-dashed lines represent intensity thresholds of 1.25×10^5^ a.u. (dark grey) and 2.25×10^5^ a.u. (light grey). For pp71^WT^, 0.25% of virions >1.25×10^5^a.u. and 0% of virions >2.25×10^5^a.u.; for pp71^HI^, 7.1 % of virions >1.25×10^5^a.u. and 1% of virions >2.25×10^5^a.u. (these percentage are shown in Fig. 3*F*). (*B*) Left panel: qRT-PCR analysis of viral genome copy number in pp71^HI^ and pp71^WT^ stocks after stocks were normalized to have equivalent infectivity by infectious units (* p < 0.05, two-tailed t test). Right panel: Infectivity of same pp71^HI^ and pp71^WT^ stocks, as measured by TCID50, after stocks were matched to have equivalent genome copy numbers (* p < 0.05, two-tailed t test). (*C*) Upper panels: Confocal micrographs of IE2-YFP reporter-virus infections (MOI 0.01) on human fibroblasts (HFFs) or a pp71-expressing HFF (WF28) (Scale Bar: 200 µM). Lower panels: Flow cytometry analysis of lytic IE2-YFP expression on indicated days. (*D*) Viral replication (titer) quantified by TCID50 seven days post infection of HFFs or pp71-expressing HFFs. (*E*) Schematic of ‘Reversion assay’ to test if pp71^HI^ virus harbors secondary mutations that influence its phenotype. If pp71^HI^ phenotype has a genetic component, lower genome-copy number (i.e., higher infectious particle-to-genome ratio) relative to pp71^WT^ will be retained and selected for due to its replicative advantage. In contrast, if the pp71^HI^ phenotype is non-genetic, excess packaged pp71 will not be retained on low passage in naïve cells and phenotype will revert to pp71^WT^ (i.e., genome-copy number and infectious particle-to-genome ratio equal to pp71^WT^). (*F*) qPCR analysis of virus output from the ‘reversion assay’ (i.e., pp71^HI^ virus after low passage in HFF). Titer of the resulting virus was measured by TCID-50, matched to wild-type (as in Fig. 2*B*), and viral genomes in the MOI-matched isolate then quantified by qPCR (difference is not significant by t-test).

We tested the functional consequence of increased pp71 by comparing the number of viral genomes present in pp71^WT^ and pp71^HI^ virus isolates at equivalent infectivity by qPCR. Surprisingly, three times as many viral genomes were required in a pp71^WT^ virion isolate to generate the same infectivity as a pp71^HI^ virion isolate (**Fig. 2*B***, left panel). Conversely, when the viral isolates were then equalized for the number of viral genomes, the pp71^HI^ virus was an order of magnitude more infectious than pp71^WT^ virus (**Fig. 2*B***, right panel).

Next, we tested how pp71 levels influence multi-round viral replication kinetics by assaying expression of the lytic master regulator, the 86-kDa immediate early 2 (IE2) protein. We passaged an IE2-YFP expressing virus (31) on cells that conditionally overexpressed pp71 during late infection to produce virions carrying excess pp71 (32) (i.e., produce pp71^HI^ virus). As predicted (31), increased pp71 generated increased single-round viral production as measured by IE2 expression (**Fig. 2*C***), and passaging virus on conditionally pp71-overexpressing cells, to continually produce pp71^HI^ virus, generated a >1-log increase in viral titer (**Fig. 2*D***). The increased number of IE-expressing cells on day 5 (i.e., ∼24 hours into the second round of infection) could not be accounted for by ectopic cellular pp71 in the overexpression cell line, since pp71 expression in this cell line is driven by a promoter activated late in CMV infection, and no increased IE expression was observed at 12 or 24 hours post infection (SI Appendix, **Fig. S3**), in agreement with previous reports (29).

To verify that the pp71^HI^ phenotype was not a result of a secondary mutation in the virus, we performed a ‘reversion assay,’ in which pp71^HI^ virus was minimally passaged in fibroblasts that did not overexpress pp71 (**Fig. 2*E***). We hypothesized that if a genetic mutation in the virus was responsible for the pp71^HI^ phenotype, the infectivity of the resulting ‘output’ virus would remain higher than pp71^WT^ (i.e., the particle-to-genome ratio would remain three times higher than pp71^WT^). Furthermore, given the increased replicative fitness of pp71^HI^, any underlying mutation should be selected for, undergo ‘fixation’, and be maintained indefinitely. In contrast, if the phenotype was non-genetic and solely due to excess packaging of pp71, the excess pp71 would be diluted out during low passage, and the infectivity of the resulting ‘output’ virus would revert to pp71^WT^. qPCR analysis indicated that the number of viral genomes present in MOI-matched pp71^WT^ and ‘output’ isolates was roughly equivalent (**Fig. 2*F***), supporting a non-genetic basis for the pp71^HI^ phenotype.

### High pp71 carries an evolutionary cost by reducing latency establishment

The enhanced infectiousness (**Fig. 2*B***) and replicative fitness (**Fig. 2*C***) conferred by higher pp71 levels presented an evolutionary conundrum given the broad virion-to-virion pp71 distribution (**Fig. 1*B***). Specifically, Darwinian selection theory predicts that, absent a counterbalancing selection pressure, virus variants that package higher pp71 levels should be selected for to promote replicative fitness. Thus, virus variants with high pp71 should sweep the population such that the virion-to-virion pp71 distribution would be narrow (low variability) with a high mean level, as opposed to the observed broad (high variability) pp71 distribution. To address this conundrum, we hypothesized that virions with lower pp71 levels may provide the putative counterbalancing selection pressure via an advantage in silencing lytic gene expression that is required to establish viral latency.

Previous studies demonstrated that during HCMV infection of undifferentiated cells, pp71 is excluded from the nucleus, promoting IE silencing and latency (33). Building off this finding, we hypothesized that pp71^HI^ virus may overcome the putative threshold for nuclear exclusion and allow a fraction of pp71 to penetrate the nucleus, leading to de-silencing and lytic expression. To test this, we infected embryonal undifferentiated NTera2 cells, an established latency model (34), with a newly developed two-color dual-reporter virus TB40E-IE-mCherry-EYFP (based on CMDR mutant described previously (18)). In this reporter virus, mCherry-only fluorescence corresponds to IE1-only expression, whereas YFP fluorescence indicates that IE2 is expressed. TB40E-IE-mCherry-EYFP was packaged to generate either pp71^WT^ or pp71^HI^ virus, and lytic IE expression was assayed in undifferentiated NTera2 cells (**Fig. 3*A***). No IE1 or IE2 expression was observed upon pp71^WT^ infection, whereas pp71^HI^ virus generated a significant percentage of dual-positive cells (**Fig. 3*A*–*B***, *SI Appendix*, **Fig. S4A**). Using confocal immunofluorescence microscopy, we then directly imaged the sub-cellular localization of pp71 and IE2 shortly after infection of undifferentiated NTera2 cells (**Fig. 3*C***). The pp71^HI^ virus generated a striking increase in nuclear pp71 levels compared to pp71^WT^ virus and nuclear IE2 expression levels in this latency model correlated with nuclear pp71 penetrance (**Fig. 3*D***). A reversion assay in NTera2 cells verified that this pp71^HI^ phenotype in NTera2 cells is non-genetic (*SI Appendix*, Fig. S5A−C). In addition, IE-expression generated by pp71^HI^ could be phenocopied by supplying pp71 *in trans* (i.e., transient transfection of NTera2 cells with a pp71 expression vector) and infecting with pp71^WT^ virus (*SI Appendix*, **Fig. S5D**−**E**).

**Figure 3:**
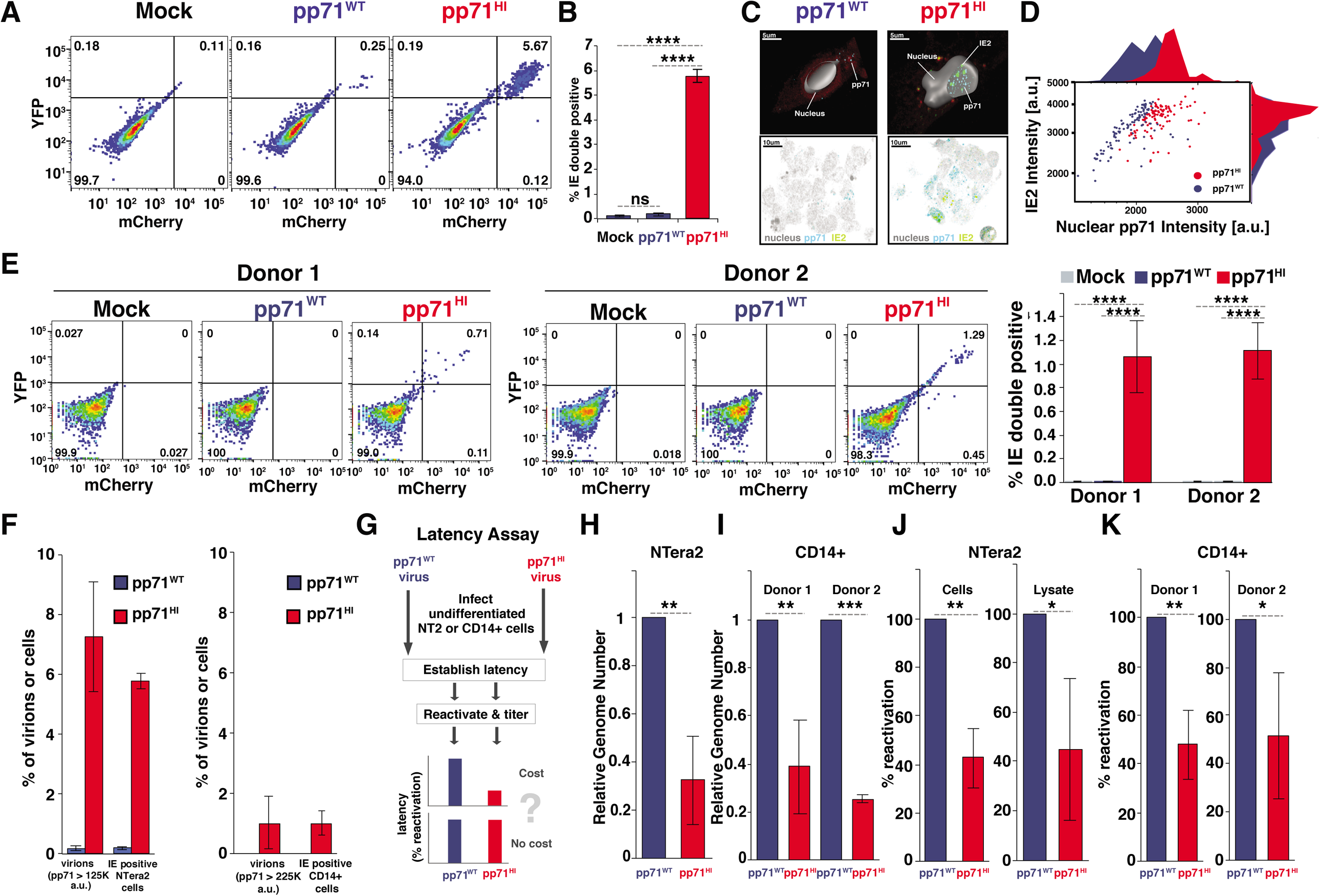
The ‘cost’ of high pp71: High virion pp71 levels overcome nuclear exclusion in undifferentiated cells and impede establishment of viral silencing. (*A*) Flow cytometry analysis of undifferentiated NTera2 cells infected with dual-reporter TB40E-IE-mCherry-EYFP. Cells were either mock infected (left), infected with pp71^WT^ virus (middle) or with pp71^HI^ virus (right) at MOI=3. (*B*) Quantification of % IE double-positive population for three biological replicates from the flow cytometry data shown in Fig. 3*A* (p-values from Student’s t test: ****<0.0001). (*C*) Representative confocal immunofluorescence micrographs of infected NTera2 cells (MOI=3) assayed 6 hours post-infection with either pp71^WT^ or pp71^HI^ virus expressing IE2-YFP. 2D images along with a single confocal plane from a 3D z-stack are shown along with a 3D reconstruction of the cell nucleus (solid). pp71 (stained via α-pp71 antibody) is visible as fluorescent puncta (teal) proximal to IE2-YFP puncta (green). Extra-nuclear fluorescence (red) is due to cytoplasmic auto-fluorescence. (*D*) Image-based quantification of pp71 and IE2 intensity levels in cells exhibiting both IE2 expression and nuclear pp71 levels for cells infected with pp71^WT^ (purple; 105 cells) or pp71^HI^ (red; 109 cells). Histograms on axes are derived from projecting the dot-plot data onto respective axis. (*E*) Left: Flow cytometry analysis of donor-derived human CD14+ primary monocytes infected with dual-reporter TB40E-IE-mCherry-EYFP (MOI = 2), 6 hours post-infection, for two donors. Cells were either mock infected, infected with pp71^WT^ virus or pp71^HI^ virus. Right: Quantification of %IE double-positive CD14(+) monocytes per donor, as assayed by flow cytometry 6 hours post-infection (p-values as in panel B). (F) Comparative analysis of percentage of virions with high pp71 levels (from Fig. 2*A*) vs. cells with IE expression (Fig. 3*C* and 3*E*). (*G*) Schematic of the latency assay for pp71^WT^ and pp71^HI^ virus. Briefly, undifferentiated NTera2 or CD14+ cells are infected with either pp71^WT^ or pp71^HI^ virus, latency established over 4−10 days, and virus then reactivated from latency. Latency is quantified by qPCR for viral genomes as well as titering on HFF after reactivation. (H) qPCR quantification of latent CMV genomes 4 days after TB40E infection of NTera2 cells. (I) qPCR quantification of latent CMV genomes 10 days after TB40E infection of CD14+ monocytes. (*J*) Analysis of latent reactivation in NTera2 cells (initial infection with pp71^WT^ or pp71^HI^ TB40E-IE-mCherry-YFP virus at MOI=3). 4 days after infection, NTera2 cells were treated with TSA for 24 hours, washed three times and serially diluted and co-cultured with HFFs for 10 days and PFU/ml then calculated by TCID-50. NTera2 cell lysate was titered in parallel for 10 days. Average of three biological replicates shown with pp71^WT^ titers normalized to 100% reactivation. (K) Analysis of latent reactivation in human primary CD14+ monocytes (initial infection with pp71^WT^ or pp71^HI^ TB40E-IE-mCherry-YFP virus at MOI=2). 10 days after infection, CD14+ cells were co-cultured with HFFs (10-fold serial dilution) supplemented with reactivation media (IL3, IL6, G-CSF, GM-CSF) for 15 days and viral titers then analyzed by TCID-50. The experiment was performed for two donors with three biological replicates and shown with pp71^WT^ titers normalized to 100% reactivation. (p-value <0.05 was considered statistically significant: *<0.05, **<0.01, ***<0.001, ****<0.0001, two-tailed t test).

To determine if high levels of pp71 also impeded silencing in human primary CD14+ monocytes—which serve as a reservoir for harboring latent HCMV (17, 35, 36)—we infected donor-derived primary human CD14+ monocytes with pp71^WT^ or pp71^HI^ dual-reporter TB40E-IE-mCherry-EYFP virus at MOI = 2, and assayed for IE expression by flow cytometry (**Fig. 3*E***, *SI Appendix*, **Fig. S4B**). In agreement with the NTera2 data, in primary monocytes from two separate donors pp71^HI^ infection generated a significant number in IE dual-positive cells, whereas pp71^WT^ virus generated no detectable IE expression (**Fig. 3*E***). Immunofluorescence imaging verified that pp71 was only present in the nucleus in CD14+ monocytes infected with pp71^HI^ virus, and correlated with IE expression (*SI Appendix*, **Fig. S5G**).

To determine if the increased pp71 in pp71^HI^ virions could explain the frequency of IE-positive undifferentiated cells, we analyzed the percentage of virions with the highest pp71 levels in both the pp71^HI^ and pp71^WT^ isolates, compared to the percentage of IE-expressing cells generated. The virion-imaging data (**Fig. 2*A***) show that ∼0.25% of pp71^WT^ virions lay above a 1.25×10^5^ a.u. intensity threshold, whereas ∼7.1% pp71^HI^ virions surpassed this threshold. In comparison, pp71^WT^ generated IE expression in 0.25% of NTera2 cells, whereas pp71^HI^ generated IE expression in 5.7% of NTera2 cells (the difference between 5.7% and 7.1% was not statistically significant) (**Fig. 3*F***). Similarly, ∼1% of pp71^HI^ virions surpassed a 2.25×10^5^ a.u. threshold, and ∼1% of CD14+ monocytes express IE after pp71^HI^ infection, whereas 0% of pp71^WT^ virions surpassed 2.25×10^5^ a.u., and 0% of monocytes express IE after pp71^WT^ infection (**Fig. 3*F***). These data indicate that virions with high pp71 abundance generate IE expression in undifferentiated cells, and likely do so by overcoming the cytoplasmic pp71 restriction.

To directly test the central tenet of bet-hedging theory—i.e., that the phenotype with a fitness advantage under one condition carries a ‘cost’ under another condition— we used an established approach (17, 34, 36) to analyze the percentage of NTera2 cells and primary CD14+ monocytes that establish and reactivate latency (**Fig. 3*G***). In this assay, cells are infected with either pp71^HI^ or pp71^WT^ virus and then incubated until IE expression silences—i.e., 4 days for NTera2 and 10 days for CD14+ cells (*SI Appendix*, **Fig. S6A**−**B**)— latency establishment is quantified by the number of genome copies, and latency reversal is quantified on permissive cells. If pp71^HI^ virus carries a ‘cost’, latency establishment and reactivation are predicted to be diminished in comparison to pp71^WT^ virus. In agreement with bet-hedging theory, pp71^HI^ virus exhibited a significant reduction in both latency establishment (**Fig. 3*H, I***) and reactivation (**Fig. 3*J, K***) in both NTera2 cells and CD14+ monocytes. Collectively, these data indicate that increasing virion-associated pp71 abundance promotes a lytic-expression program in undifferentiated cells leading to reduced latency establishment.

### Super-Poissonian variability in virion tegument abundance enables bet-hedging

Overall, our results show that heterogeneity in tegument protein abundance in CMV viral particles creates a phenotypically diverse virion population, providing a molecular mechanism for probabilistic bet hedging between lytic and latent infection (**Fig. 4**). Interestingly, a positive auto-regulatory circuit mediated by the 76-kDa immediate early 1 (IE1) protein provides a mechanism for further amplification of pp71 phenotypic diversity. Positive feedback loops are known to amplify variability and generate bimodality in other viral and non-viral systems (37, 38). We speculate that variable pp71 levels in infecting virions provide an initial ‘kernel’ level of variability, which is then processed by IE1 positive-feedback circuitry. If the incoming pp71 level in the infecting virion is *below* the threshold for activation of the IE1 positive-feedback circuit, it will undergo proteolysis within its characteristic 8-hr half-life without sufficiently activating the immediate-early viral expression program, and the infected cell will enter a quiescent state of expression. In contrast, if the incoming pp71 level in the infecting virion is *above* the threshold, the immediate-early viral expression program will be activated and the cell will proceed to lytic infection. Stochastic variability in packaged tegument levels in virions may explain the differing data on early IE expression in undifferentiated cells (36, 39, 40).

**Figure 4:**
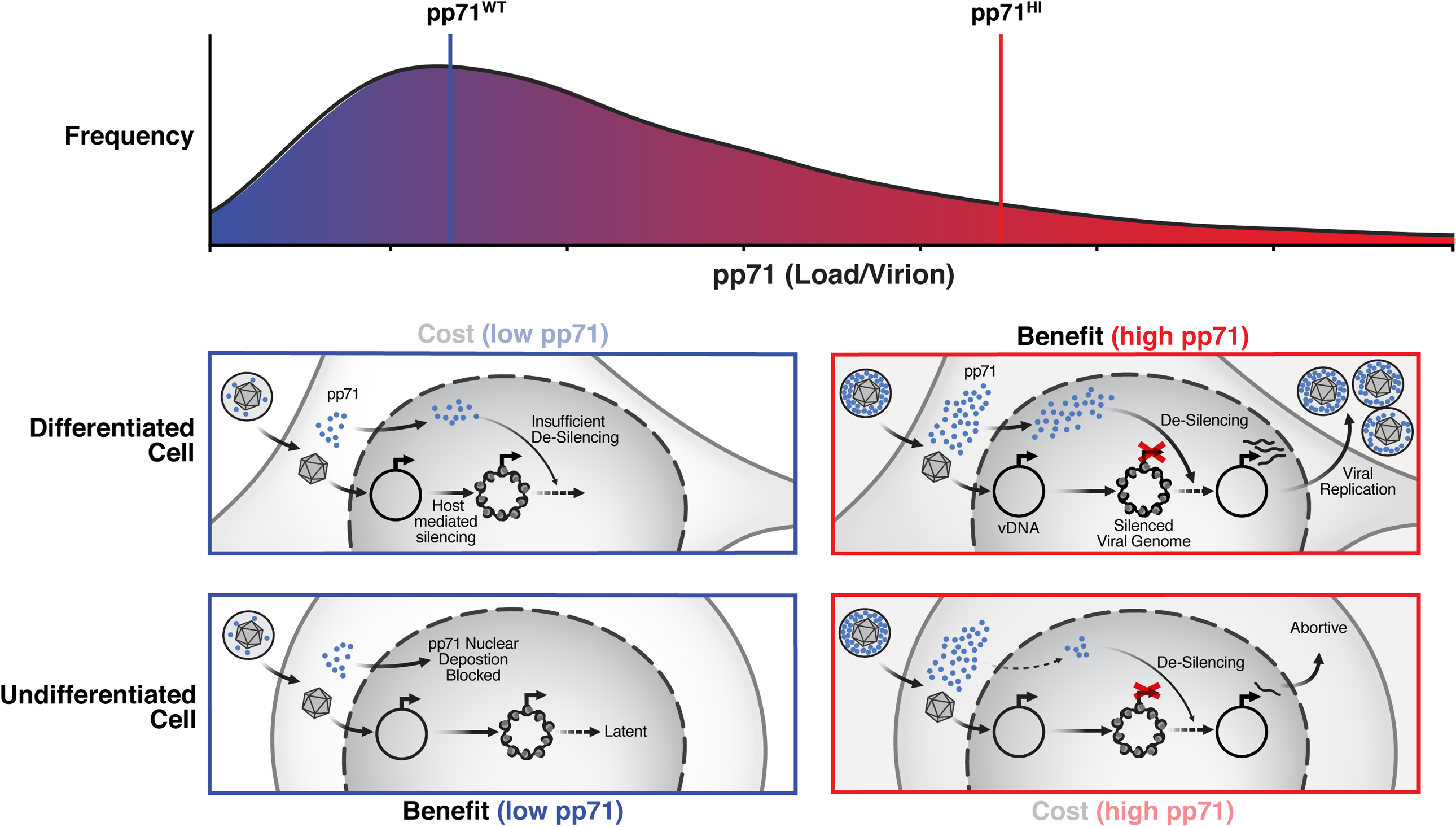
Summary model: super-Poissonian variability in tegument abundance enables bet-hedging between replication and latency. In differentiated cells (middle panels), pp71^HI^ virions (right) have an ‘advantage’ over virions with low pp71: pp71^HI^ enables de-silencing and enhanced replicative fitness compared to pp71^WT^ virions that are more likely to generate a silenced infection. In undifferentiated cells (bottom panels), pp71^HI^ virions (right) have a ‘cost’ as they impede viral silencing and reduce establishment of latency. In undifferentiated cells, virions with low pp71 levels promote establishment of viral silencing and latency.

Stochastic tegument packaging has both similarities and differences with other proposed bet-hedging mechanisms. In a sense, the stochastic tegument packaging mechanism can be considered a phenomenological analogue of early bet-hedging theories of variability in seed-husk thickness (1). In influenza virus, variations in virus composition, morphology, and sequence also promote phenotypic diversity (6, 41), whereas in the lentiviridae, probabilistic establishment of proviral latency and persistence is regulated by stochastic fluctuations in viral transcription (37, 42, 43), with the HIV promoter generating very large stochastic fluctuations in gene expression. Surprisingly, the variability in virion pp71 levels (**Fig. *1***) is extreme, even in comparison to the variability generated by HIV (44, 45), with σ/μ and σ^2^/μ^2^ values being ∼5-fold greater. As such, stochastic tegument packaging may represent a mechanism for further amplifying phenotypic variability downstream of gene expression fluctuations. Tegument protein loads also appear variable in other herpesviruses (22-24), suggesting that stochastic packaging may be a generalized mechanism for probabilistically maximizing fitness in variant host environments and immune-stimulation conditions. Overall, approaches to modulate tegument expression levels may provide an avenue for limiting viral persistence or, alternatively, reversing viral silencing and persistence.

## Abbreviations

crs: *cis* repression sequence
CMV: human cytomegalovirus
HFF cells: human foreskin fibroblast cells
MIE: Major Immediate-Early
MIEP: Major Immediate-Early promoter
MOI: multiplicity of infection
PFU: plaque-forming units
TCID_50_: 50% tissue culture infective dose
UV: ultraviolet
VPA: valproic acid
YFP: yellow fluorescent protein
DiI: 1,1’-dioctadecyl-3,3,3’3’-tetramethylindocarbocyanine perchlorate

## AUTHOR CONTRIBUTIONS

SC, NV, JK, CBF, and LSW conceived and designed the study. SC, KD, MW, JK, CBF, and NV performed the experiments. SC, MW, JK, CBF, NM, BH, and LSW analyzed the data. NM, MP, and FG provided reagents. SC, NV, JK, and LSW wrote the paper.

## COMPETING INTEREST STATEMENT

The authors declare no competing interest.

## ACKNOWLEDGEMENTS

We thank Edward Mocarski for generously providing us with pp150 expression plasmid and Thomas Stamminger for the generous contribution of the pp71-YFP reporter virus. We thank Elena Ingerman, Melanie Ott, JJ Miranda, Marielle Cavrois, Nandhini Raman, Elizabeth Tanner, Nadia Roan, Jason Neidleman and the Weinberger lab for discussions and suggestions, Melissa Teng and Jinny Wong for technical support, and Kathryn Claiborn for reviewing the manuscript. We acknowledge the Gladstone Flow Cytometry Core, funded through NIH P30 AI027763. SC was supported in part by generous donations to the Cynthia Bolovan-Fritts Memorial Fund. We acknowledge M. Ghassemian at the Biomolecular and Proteomic Mass Spectrometry Facility at UCSD for SILAC mass spectrometry technical support (funded through NIH S10 OD016234 and S10 OD021724). LSW acknowledges support from the Bowes Distinguished Professorship, the Alfred P. Sloan Research Fellowship, and the NIH Director’s New Innovator Award (OD006677) and Pioneer Award (OD17181) programs

## MATERIALS AND METHODS

### Virus cloning and purification

Viral recombinants TB40E IE2-YFP, HCMV AD169 pp71-EYFP, pp150-EYFP, dual-tagged TB40E-IE-mCherry-EYFP have previously been described (18, 30, 31, 46, 47)—the two-color TB40E-IE-mCherry-EYFP dual-reporter virus was cloned as an intermediate of the previously described CMDR virus (18). Viruses were propagated in MRC5 human foreskin fibroblast (HFF) cells (American Type Culture Collection) and upon infection reaching ∼90% viral cytopathic effect or ∼90% GFP, the culture supernatant was collected and filtered through a 0.2 μm filter. HCMV pp71HI virus was generated by expanding either HCMV AD169 pp71-EYFP virus (30) or IE2-YFP virus (31) on life-extended HFFs (21) stably transduced with either a pp71-EYFP or a pp71 expression vector, respectively, as described (30).

To purify particles for single-virion imaging, viral recombinants (HCMV AD169 pp71-EYFP or pp150-EYFP, and HSV-1 Strain F VP26-EGFP) were used to synchronously infect 10 confluent 15cm culture dishes containing either MRC5 cells (HCMV AD169) or Vero cells (HSV-1 Strain F). Supernatants were collected from infected tissue-culture plates after a single round of replication, corresponding to approximately 30 h.p.i. (hours post infection) for HSV-1, 96 h.p.i. for HCMV AD169 pp71-EYFP, and 168 h.p.i. for HCMV AD169 pp150-EYFP. Supernatants were then clarified by low speed spin at 3000 rpm for 10 minutes and the resulting debris pellet was discarded. For HCMV purifications, the clarified supernatant was then layered over a 20% sorbitol gradient supplemented with 100 ug/mL Bacitracin. For HSV-1 purifications, supernatant was layered over a 30% sucrose gradient supplemented with 100 ug/mL Bacitracin. All overlays were centrifuged in a Beckman Coulter Optima L-80 XP Ultracentrifuge in an SW-28 rotor at 20k rpm and 18°C for 1.5 hours. Pellets were re-suspended in 1 mL of TN buffer and overlaid on top of a glycerol tartrate gradient (48). This gradient was then centrifuged in a Beckman Coulter Optima L-80 XP Ultracentrifuge in an SW-41 rotor at 28k rpm and 4°C for approximately 15 minutes. Lower bands corresponding to the infectious virion population were harvested by sequentially removing upper layers of the gradient and isolating the appropriate band. DiI was then added to isolated bands such that the final concentration was 10% v/v and allowed to incubate for 2 hours protected from light at room temperature. After incubation, the samples were fixed in 2% (v/v) formaldehyde (Tousimis Research Corporation, cat# 1008A diluted 1:10 in PBS).

### Super-resolution imaging of viral particles

All viral purifications were mounted on 3” x 1” x 1 mm microscope slides (Fisherfinest, #12-544-1) with plasma-treated 22mm x 22mm #1.5 coverslips (Fisherbrand, #12-541-B). A Plasma Etch PE50 unit was used to clean coverslips at 100W for 5 minutes prior to mounting. 3 uL of each purification was dotted onto the microscope slide and overlaid with a plasma-treated #1.5 coverslip. Coverslips were sealed with fast-drying clear nail polish. All mounted slides were stored at 4°C and protected from light until imaging. Viral particle purifications were imaged at super resolution using a Zeiss LSM880 Airyscan microscope using a 63x 1.4NA oil-immersion DIC M27 objective (Apochromat) and laser lines for 488 nm and 561 nm. Each x-y position was sampled at 10 z planes (1.665 μm intervals) to identify the z plane where virions particles were at maximal concentration and this z was chosen for image analysis. For confocal imaging of the viral particles, a Zeiss Observer Z1 Yokagawa Spinning Disk Confocal Microscope equipped with a Yokogawa spinning disk, a CoolSNAP HQ2 14-bit camera (PhotoMetrics, Tucson, AZ), and laser lines for 488nm and 561nm. was used with a 100x 1.45 NA oil-immersion objective (Apocromat). Zeiss Immersol 518F immersion oil was used during imaging of all purifications (Carl Zeiss Microscopy, #444970-9000-000). Viral particle preparations were imaged under a 488 nm laser excitation (3.5s exposure) at 20% laser power and a 531 nm laser for 300ms at 11.5% laser power.

### Super-resolution image analysis

Super-resolution images of purified particles were analyzed using an established superresolution algorithm (26) that restricted analysis to particles 100nm−300nm in diameter and fit each particle’s intensity distribution to a single Gaussian distribution to exclude doublets. Mean intensity of each particle was calculated from the fitted single-Gaussian distribution. Point-spread functions were calculated and images were also size-standardized against fluorescently-labeled size-calibration microspheres (TetraSpeck™ ThermoFisher) imaged under the same conditions. For parallel confocal imaging of purified viral particles, images were analyzed using a custom MATLAB script. Briefly, the algorithm is designed to first locate and separate potential virion particles from background pixels in a micrograph. Once thresholded, each of the pixels in a segmented particle was normalized to the maximum pixel intensity within that same segmented particle. An additional normalized threshold for pixel intensity of 50% was introduced in order to isolate and characterize the peak of the PSF associated with each particle (akin to Full-Width at Half-Maximum (FWHM)). Once segmented and normalized, several quality control steps were implemented in order to remove objects unlikely to be single viral particles. Firstly, a strict pixel-area size threshold of 4-11 pixels was introduced for segmented objects based upon the measured camera pixel scaling of .129 um per pixel. These pixel area values were verified against size-standardized fluorescently-labeled microspheres imaged under the same conditions. Following size exclusion, a series of exclusion steps based on the composition and shape of segmented particles was used. The final quality control step was achieved by identifying objects with at least 1-pixel intersections between the green and red channels. These co-localized particles were then quantified by summing pixel intensities within a co-localized viral particle to produce a cumulative sum of fluorescence intensity for an individual virion. Quantifications of viral particles from each set of micrographs were then pooled together and placed into corresponding histograms. Virion particles with fluorescence intensities +/- 1 standard deviation from their respective means were excluded to remove any possible contribution from viral dense bodies or twinned viral particles.

### Cell-culture conditions and drug perturbations

Human foreskin fibroblasts (HFF), MRC5 fibroblasts, and telomerase life-extended HFFs (21) were maintained in Dulbecco’s Modified Eagle’s Medium (DMEM) supplemented with 10% fetal bovine serum (FBS) and 50 U/ml penicillin-streptomycin at 37°C and 5% CO_2_ in a humidified incubator. NTera2 cells were obtained from ATCC (American Type Culture Collection) and maintained in DMEM supplemented with 10% FBS and 50 U/ml penicillin-streptomycin at 37°C and 5% CO_2_ in a humidified incubator.

### Negative staining and electron microscopy

Infectious virion was purified using density gradient centrifugation as mentioned above and negative staining of virion was performed. Briefly, virion was diluted 1:100 in PBS buffer, 2uL of virion was added on holey film TEM grid (EM Resolutions, Sheffield, United Kingdom), washed three times with distilled water and stained with 10ul of 2% phosphotungstic acid for 1min. Grids were subjected to electron microscopy using JEOL JEM-1230 transmission electron microscope (JEOL, USA).

### Viral titering and qPCR analysis of viral genome copy numbers

pp71^WT^ and pp71^HI^ viruses were generated by expanding TB40E IE2-YFP or TB40E-IE-mCherry-EYFP virus on HFF or WF28 HFF cells that were stably transduced with a pp71 expression cassette (30), respectively. Viral preps were titered by TCID-50. For analysis of viral genome copy numbers, viral DNA was extracted from equal amounts of virus using a Nucleospin virus kit (MACHEREY-NAGEL, 740983.10). To account for potential differences in efficiency of DNA extraction, 5ng of a reference plasmid (pNL4-3, Addgene) was spiked in to each DNA preparation and used for qPCR normalization (Table S1). The number of viral genomes was normalized to the reference carrier DNA level in each sample. The relative difference in genomes between the two viral preparations was then used to match the viral stocks by viral genome copy number.

### Confocal imaging and flow cytometry

Confocal imaging of IE2-YFP, IE-mCherry-EYFP, and α-pp71 stained NTera2 cells was performed on an Axiovert inverted fluorescence microscope (Carl Zeiss, Oberkochen, Germany), equipped with a Yokogawa spinning disk, a CoolSNAP HQ2 14-bit camera (PhotoMetrics, Tucson, AZ), and laser lines for 488nm and 561nm. Flow cytometry analysis of IE2-YFP, and IE-mCherry-EYFP expression for HFF, NTera2, and human primary CD14+ monocytes was performed with on an LSRII flow cytometer (BD Biosciences).

### Donor-derived human peripheral blood monocyte isolation

Human peripheral PBMCs were isolated from de-identified blood samples obtained from the University Medical Center at the University of Arizona, in accordance with the Institutional Review Board. Briefly, 240mls of blood samples were centrifuged through a Ficoll Histopaque 1077 gradient (Sigma-Aldrich, St. Louis, MO) at 200 ×g for 30 min at room temperature (RT). Mononuclear cells were collected and washed six times with sterile 0.9% sodium chloride saline solution (Baxter) to remove platelets at 200 × g for 10 min at RT. Monocytes were then layered on top of a 45% and 52.5% isosmotic Percoll gradient and centrifuged for 30 min at 400 × g at RT yielding an average of 90% monocyte population.

Cells were washed three times with saline at 200 × g for 10 min at RT to remove residual Percoll and suspended in RPMI 1640 medium (Cellgro, Manassas, VA) supplemented with 5% human serum (Sigma-Aldrich, St. Louis, MO), unless otherwise stated. University of Arizona Institutional Review Board and Health Insurance Portability and Accountability Act guidelines for the use of human subjects were followed for all experimental protocols in our study.

### Virus Reactivation Assay

Virus reactivation assay was performed for two different cell types. Virus reactivation assay in Ntera2 cells was performed as described previously(34) with minor modifications. Briefly, NTera2 cells were infected with pp71^WT^ or pp71^HI^ dual-reporter TB40E-IE-mCherry-EYFP virus at an MOI=3, and at 4dpi, cells were treated with TSA (100ng/ml) (Sigma-Aldrich, St. Louis, MO) and incubated for 24h followed by washing cells three times in PBS and co-culturing at ten-fold dilution with HFFs in a 96 well plate or performing TCID50 with the cell lysate. Cells were scored for YFP and mCherry at 15 days post serial dilution. To make sure there was no reactivation of virus prior to TSA treatment, cells were lysed at 4dpi and the lysate was added to HFFs and monitored for IE expression for 15 days (*SI Appendix*, Fig.S7A). Virus reactivation assay in CD14+ cells was performed as described previously with minor modifications(17). Briefly, for CD14+ cells, cells from two donors were infected with pp71^WT^ or pp71^HI^ dual-reporter TB40E-IE-mCherry-EYFP virus at an MOI=2, at 10dpi when the virus has established latency, cells were 2-fold serially diluted and co-cultured with fibroblast in a 96-well plate with reactivation media (RPMI supplemented with 20%FBS, 100u/ml penicillin, 100mg/ml streptomycin, 20ng/ml each IL3 IL6, G-CSF, GM-CSF (Sigma-Aldrich, St. Louis, MO). Fibroblasts were monitored for YFP, mCherry expression for 15 days. Equal number of cells were lysed, serially diluted, and plated on HFFs as a control (*SI Appendix*, Fig. S7B). The frequency of reactivation was calculated by TCID50 assay and %reactivation for pp71^HI^ was reported relative to %reactivation for pp71^WT^ virus, which was normalized to 100%. To quantify viral genome copy number total DNA was extracted from cells and subjected to qPCR on 7900HT Fast Real-Time PCR System (catalog no: 4329003, Thermo-Fisher Scientific) using Fast SYBR Green Master Mix (catalog no: 4385612, Applied Biosystems) and normalized with β-actin using sequence specific primers (Table S1).

### Immunofluorescent microscopy, transient transfection and Western blots

For immunofluorescent microscopy, NTera2 cells were grown on glass coverslips and infected with either pp71^WT^ or pp71^HI^ dual-reporter TB40E-IE-mCherry-EYFP virus (MOI=3) for 6h, coverslips were harvested and washed with phosphate-buffer saline (1X) (PBS, Sigma-Aldrich, St. Louis, MO) three times, and fixed, permeabilized and immunostained as described below. CD14+ cells (grown in suspension) were infected with either pp71^WT^ or pp71^HI^ dual-reporter TB40E-IE-mCherry, EYFP (MOI=2). At 6h post infection, cells were harvested by centrifuging at 900RPM, washed three times in PBS, fixed, permeabilized and immunostained as mentioned below. Briefly, cells on glass coverslips (NTera2) or in suspension (CD14+) were fixed using 4% paraformaldehyde (pH7.4)

(Electron Microscopy Sciences, Hatfield, PA) for 10 minutes on ice, washed three times with 500mL PBS, permeabilized using 0.1% TritonX-100 in PBS for 10 minutes, and immunostained. Cells were incubated with primary antibodies (pp71 antibody kindly provided by Thomas Shenk, dilution-1:100 in 0.1% BSA; Daxx antibody (MABE1911, Sigma-Aldrich, St. Louis, MO), dilution-1:50 in 0.1% BSA) for 3h at room temperature with gentle shaking followed by three PBS washes. Cells were incubated with secondary antibody (goat anti-mouse Alexa fluor 405, catalog no: A-31553 (Invitrogen™, Thermo Fisher Scientific), dilution 1:500 or donkey anti-mouse Alexa fluor 647, catalog no: A-31571(Invitrogen™, Thermo Fisher Scientific), dilution: 1:500 in PBS supplemented with 0.1% BSA) for 1h at room temperature, and washed three times with PBS. For samples stained for DAPI, cells were stained with DAPI (0.1mg/ml in PBS) at room temperature for 10 min and imaged. Transient transfection of pp150 in HFF was performed by transfecting pp150 expressing plasmid (kindly provided by Edward Mocarski) using Amaxa Nucleofection kit V (Lonza inc.). The pp71-expressing plasmid (pCGN-pp71) was transfected in NTera2 cells using lipofectamine-3000 transfection reagent (Invitrogen™) following the manufacturer’s instructions. For Western blot analysis, infectious virus was resuspended in 20 μl PBS buffer and analyzed by western blot. Briefly, 5 μl of purified virion sample was added to 1x loading buffer (100mM tris-HCl (pH6.8), 200mM DTT, 4% SDS, 0.1% Bromophenol blue, 20% glycerol), and boiled for 10 minutes at 95°C. Samples were loaded on 12% SDS-Polyacrylamide gel (Bio-Rad inc.), and ran at 90V for 2 hours in Tris-glycine running buffer (25mM Tris, 250mM glycine and 0.1% SDS). Gel was blotted on a PVDF membrane (Bio-Rad inc.) using Trans-Blot SD Semi-Dry Transfer Cell (Bio-Rad inc.) at 20V for 45 minutes, and was blocked with 10ml of Li-Cor odyssey blocking buffer (Li-Cor Biosciences Inc.) for 1 hour at room temperature with gentle shaking. Membrane was treated with primary antibody, anti-pp150 (1:200 dilution, mouse MAb 36-10, gift from Dr. William Britt) for 2 hours at room temperature with gentle shaking. The membrane was then washed three times with wash buffer (1x PBS+0.01% Tween-20), treated with secondary Li-Cor detection antibody (Li-Cor Biosciences inc.) (1:20,000 dilution, goat anti-mouse 800CW), and incubated in dark for 1 hour at room temperature. The membrane was washed three times with wash buffer and scanned on Li-Cor Odyssey system (Li-Cor Biosciences inc.).

### Data Availability

All the data is made available in the manuscript and in the *SI Appendix*.

## Notes

### Competing Interest Statement

The authors have declared no competing interest.

