## Supplementary material for "A Molecular Mechanism for Probabilistic Bet-hedging and its Role in Viral Latency": SI Appendix Figs.1-7 and Table 1

**SUPPLEMENTAL FIGURES & LEGENDS**

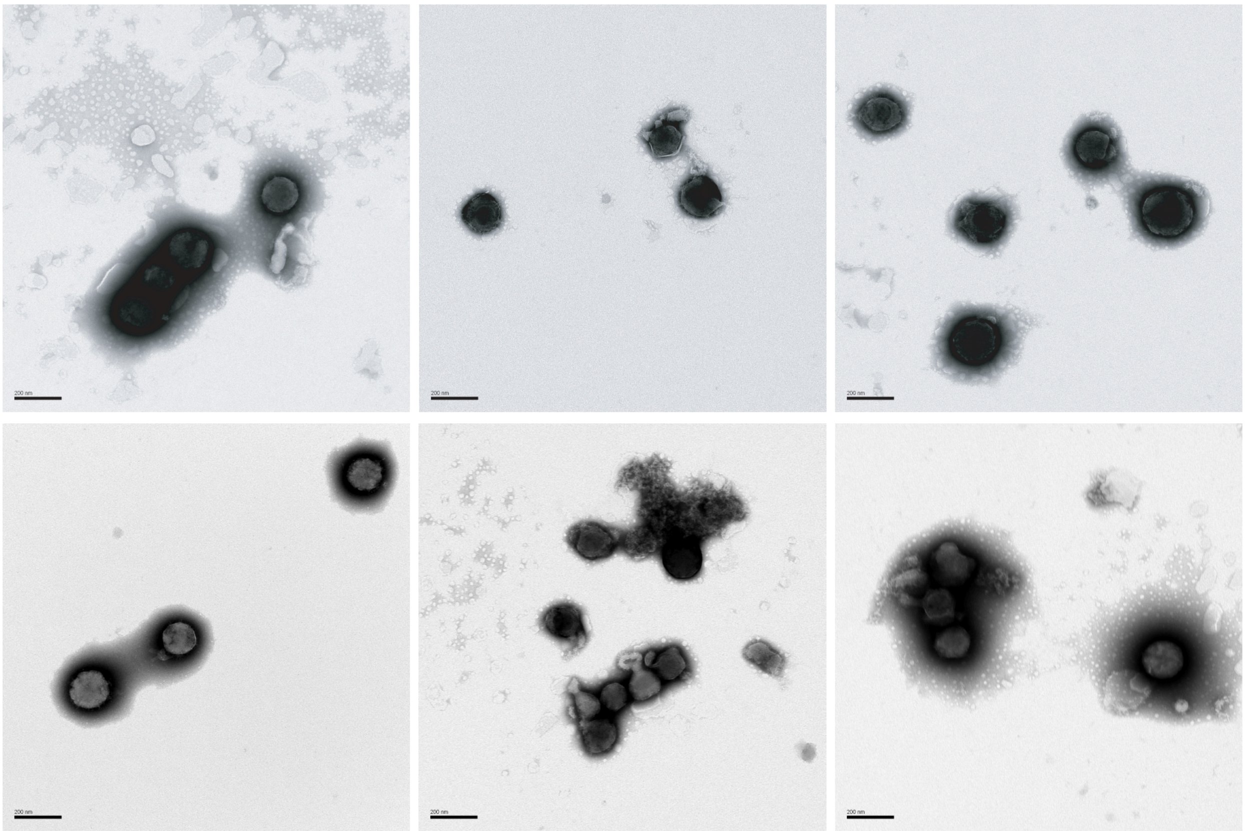

**<u>Figure S1</u>: *Electron microscopy images of purified HCMV virions*.** Representative TEM micrographs of the viral recombinant purification verifies the presence of single, structurally intact virions present after gradient purification. Scale bar = 200nm.

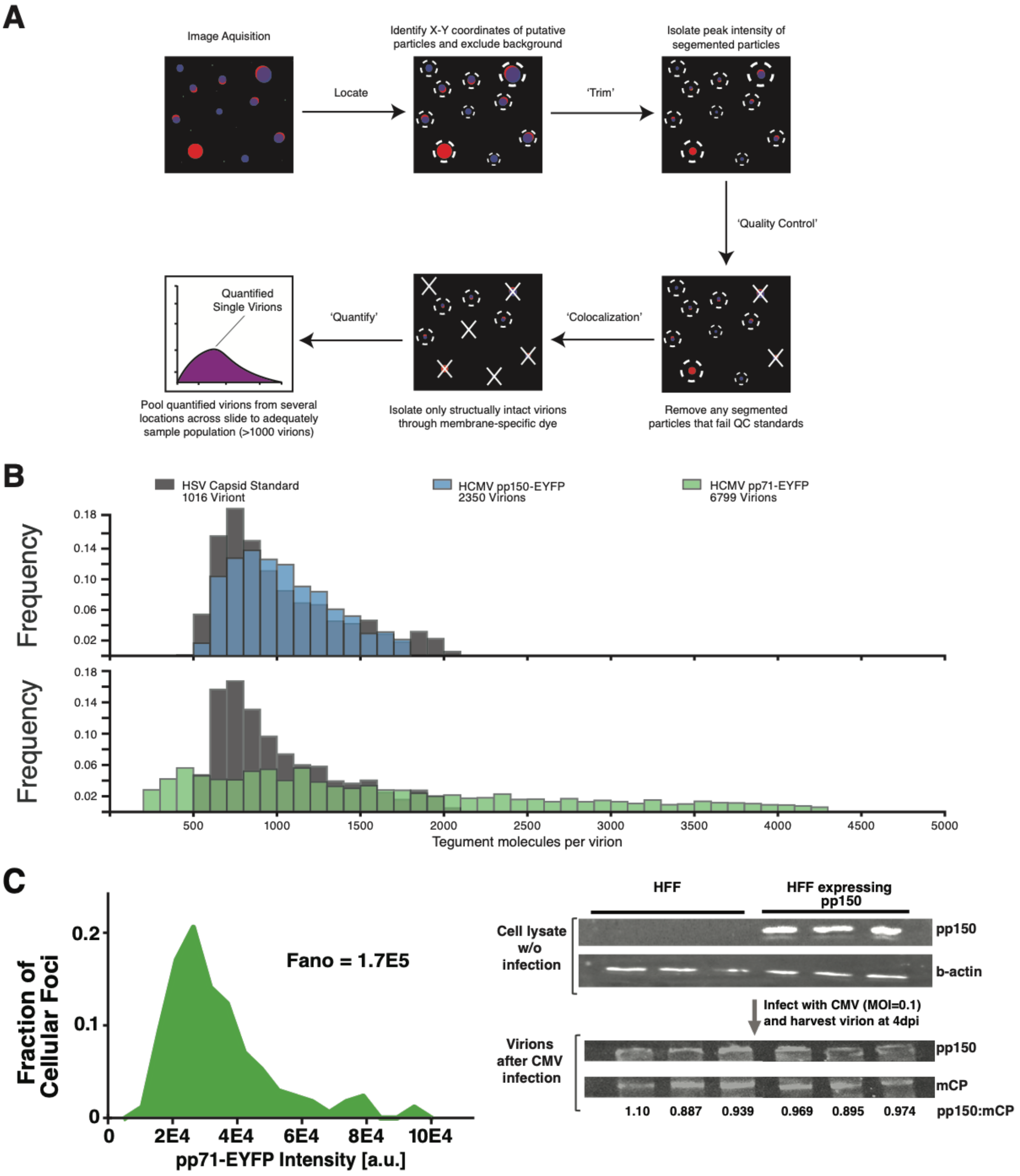
A
Image Aquisition
Locate
Identify X-Y coordinates of putative particles and exclude background
'Trim'
Isolate peak intensity of segemented particles
'Quality Control'
Remove any segmented particles that fail QC standards
'Colocalization'
Isolate only structually intact virions through membrane-specific dye
'Quantify'
Quantified Single Virions
Pool quantified virions from several locations across slide to adequately sample population (>1000 virions)
B
HSV Capsid Standard 1016 Viriont
HCMV pp150-EYFP 2350 Virions
HCMV pp71-EYFP 6799 Virions
Frequency
Tegument molecules per virion
C
Fraction of Cellular Foci
Fano = 1.7E5
pp71-EYFP Intensity [a.u.]
HFF
HFF expressing pp150
Cell lysate w/o infection
pp150
b-actin
Infect with CMV (MOI=0.1) and harvest virion at 4dpi
Virions after CMV infection
mCP
1.10 0.887 0.939 0.969 0.895 0.974
pp150:mCP

**Figure S2**: ***Substantial virion-to-virion heterogeneity in the levels of the HCMV major tegument transactivator protein pp71.*** (A) Schematic of image analysis pipeline used for quantification of tegument proteins in virions using dual-labeled viral particles. Tegument proteins were tagged with YFP while the viral envelopes were stained with DiI. Only intact viral particles in which YFP and DiI signals were co-localized were analyzed. (B) Quantification of HCMV pp71, HCMV pp150, and HSV-1 VP26 tegument copy numbers in single virion particles was performed using a custom MATLAB script. Quantification of differential variability in tegument copy numbers between virions using Fano Factor normalized to HSV-1 VP26-GFP levels. (C) (Left) Distribution of virion-sized pp71-EYFP foci intensity in an infected human fibroblast (MOI = 0.2). Nuclear pp71-YFP and pp71-YFP foci that are larger than 300nm were excluded from the analysis to analyze a total of 174 pp71-YFP particles. Also see Fig. S2. (Right) Quantification of pp150 in virions packaged in pp150 overexpressing cell-line. CMV (MOI=0.1) was grown in either control HFFs or HFFs over-expressing pp150. After 4 days, virus was extracted and subjected to sucrose density gradient centrifugation. Western blot was performed using antibodies against pp150 and UL85 (minor capsid protein), band intensity was quantified in ImageJ.

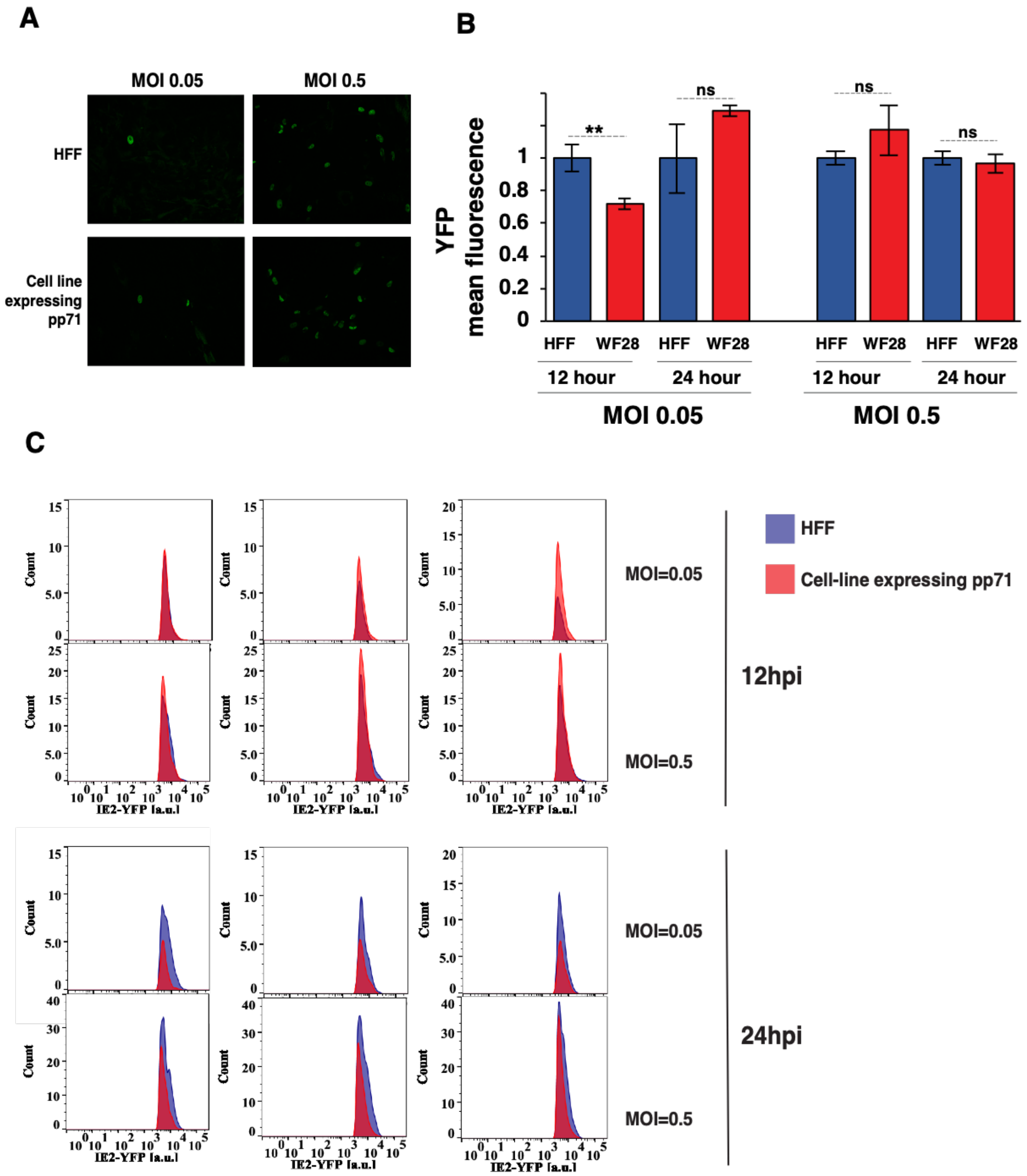


**<u>Figure S3</u>**: ***Lower frequency of lytic expression in pp71$^{WT}$ compared to pp71$^{HI}$ infected cells***. (A) Fluorescent micrographs of lytic IE2-YFP reporter virus (MOI=0.05 and MOI=0.5) in human fibroblasts or pp71 expressing pp71 HFF cell line at 24 hpi. (B, C) Flow cytometry analysis of lytic IE2-YFP reporter virus infection (MOI=0.05 and MOI=0.5) in human fibroblasts (HFFs) or pp71-expressing HFF (WF28) at 12 hour and 24 hour post infection. Quantification of IE2-YFP fluorescence (geometric mean) (B) from histograms of YFP+ population (C). bar graph and histogram color: HFF (purple), cell line expressing pp71(red).

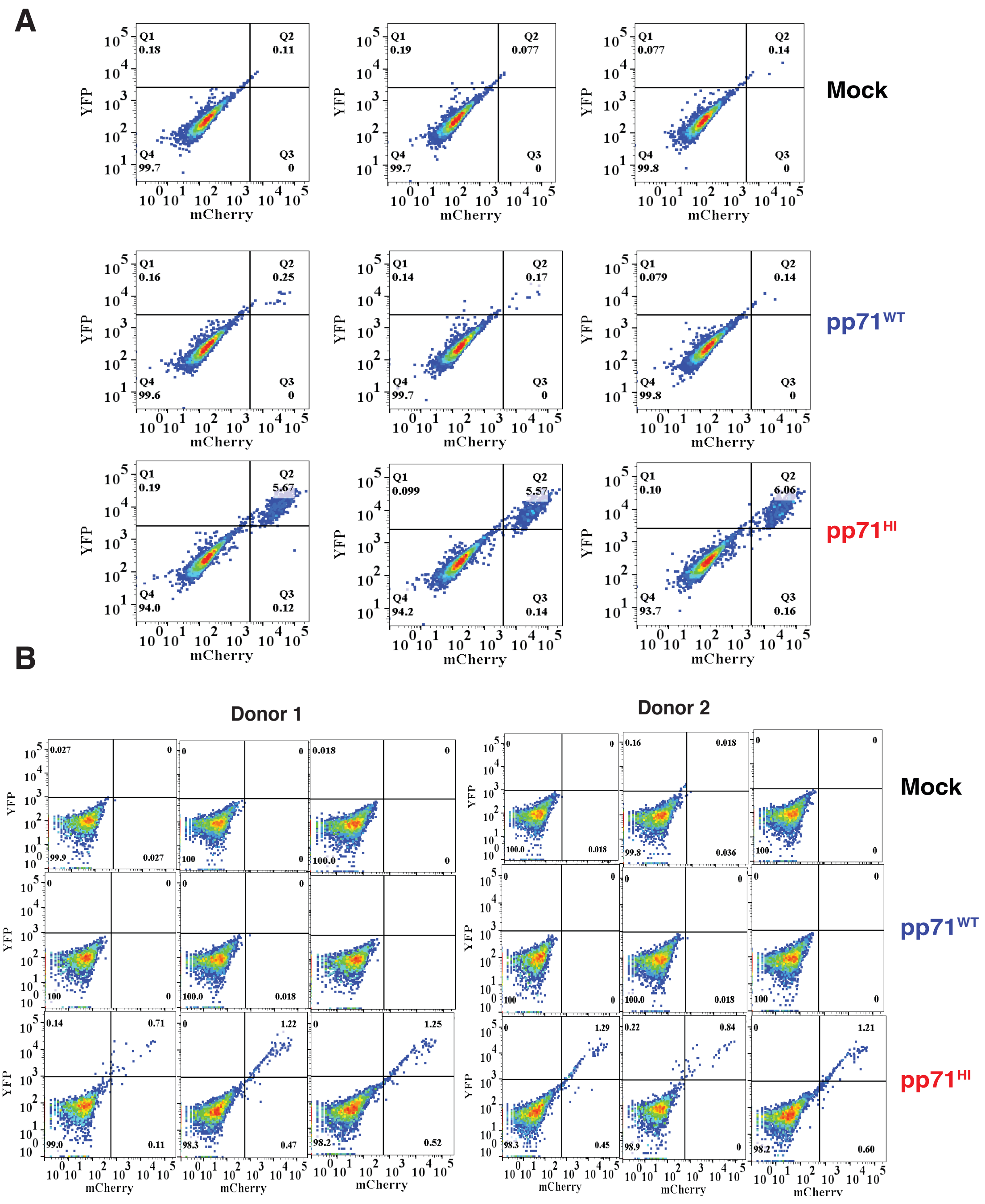


**Figure S4**. ***Replicates of infections of NTera2 and CD14+ with dual-reporter TB40E-IE-mCherry-EYFP.*** (A) Flow cytometry analysis of three biological replicates of undifferentiated NTera2 cells infected with $pp71^{WT}$ or $pp71^{HI}$ TB40E-IE-mCherry-YFP virus (MOI=3) at 6 hours post infection. (B) Flow cytometry analysis of three biological replicates of undifferentiated primary human CD14+ monocytes cells infected with $pp71^{WT}$ or $pp71^{HI}$ TB40E-IE-mCherry-YFP virus (MOI=2) at 6 hours post infection.

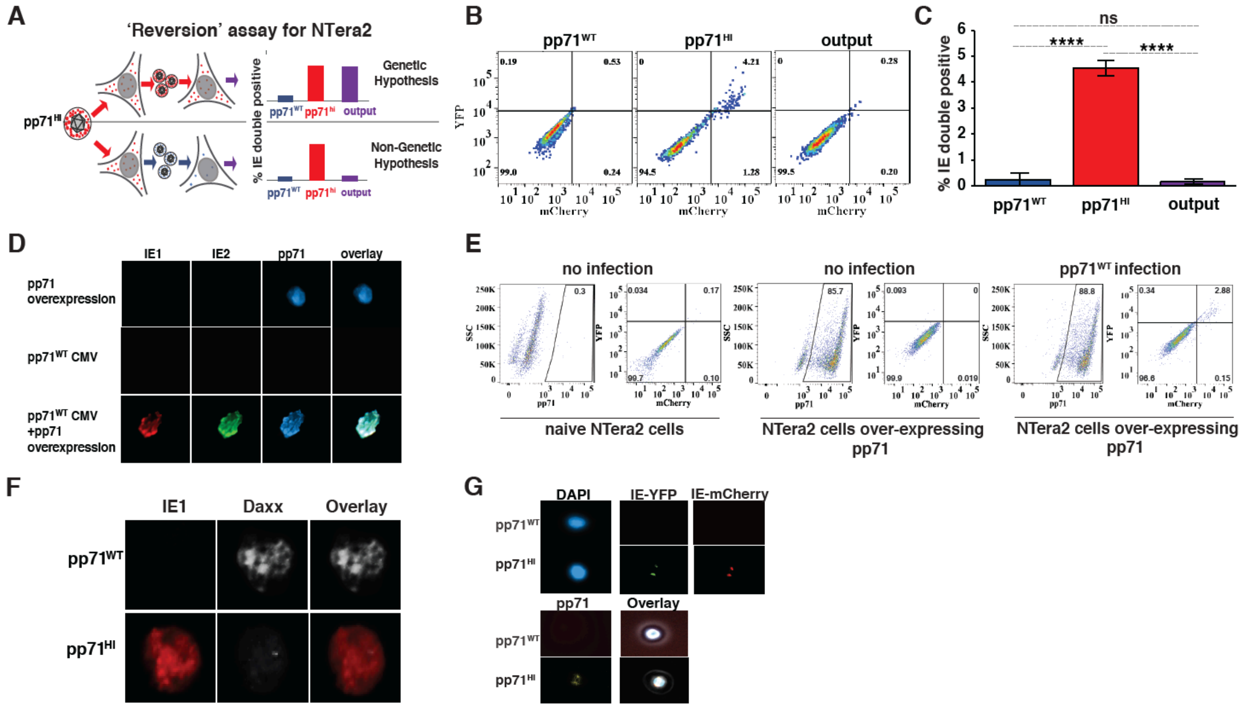


**Figure S5**: (A) Schematic of 'Reversion assay' for NTera2 cells. If pp71$^{HI}$ phenotype has a genetic component, pp71$^{HI}$ phenotype will be retained upon low passage, and 'output' virus will increase in IE expression in NTera2 cells. In contrast, if the pp71$^{HI}$ phenotype is non-genetic, excess packaged pp71 will not be retained on low passage in naïve cells and phenotype will revert to pp71$^{WT}$ (i.e., will demonstrate low or no IE expression). (B) Flow-cytometry analysis of NTera2 cells infected with pp71$^{WT}$, pp71$^{HI}$ and 'output' dual tagged TB40E-IE-mCherry virus (MOI=3) at 6hpi. (C) Quantification of % IE double positive cells from flow-cytometry analysis for NTera2 cells infected pp71$^{WT}$, pp71$^{HI}$ and output (MOI=3) at 6hpi (from Fig. S5B). (D) Ntera2 cells overexpressing pp71 were infected with pp71$^{WT}$ dual tagged TB40E-IE-mCherry virus (MOI=3), fixed at 6hpi, stained for pp71 and imaged for red=IE1, green=IE2, blue=pp71. Controls: pp71 overexpressing in Ntera2 cells by itself and pp71$^{WT}$ dual tagged TB40E-IE-mCherry CMV infection in naïve Ntera2 cells (MOI=3). (E) Quantification of pp71 and IE double positive cells from flow-cytometry data presented in Figure S5D. (F) Immunofluorescent microscopy of Ntera2 cells infected with pp71$^{WT}$ or pp71$^{HI}$ virus (MOI=3), fixed at 6hpi, and stained for Daxx, and imaged, red=IE1, white=Daxx. (G) Immunofluorescent microscopy of CD14+ cells infected with either the pp71$^{WT}$ or pp71$^{HI}$ dual tagged TB40E-IE-mCherry-YFP virus (MOI=2) at 6hpi, blue = DAPI, green = YFP, red = mCherry, yellow= pp71.

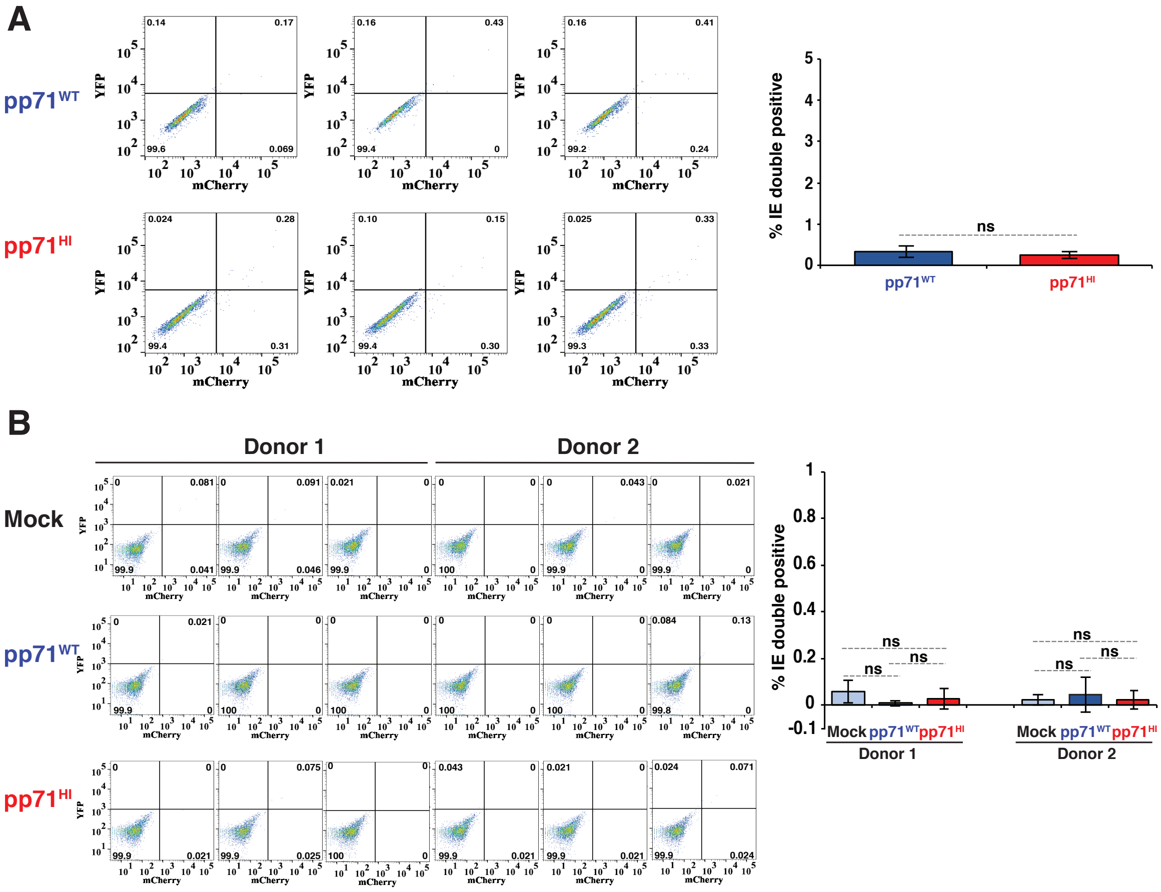


**<u>Figure S6:</u>** ***Establishment of CMV latency in NTera2 and CD14+ cells.*** (A) Flow-cytometry analysis of three biological replicates of NTera2 cells infected with pp71$^{WT}$ and pp71$^{HI}$ at 4 dpi (left), and quantification of %IE double positive at 4dpi from the flow-cytometry data (right). (B) Flow-cytometry analysis of three biological replicates of CD14+ cells from two different donors for mock, pp71$^{WT}$, pp71$^{HI}$ at 10dpi (left), and quantification of %IE double positive at 10dpi from flow-cytometry data (right).

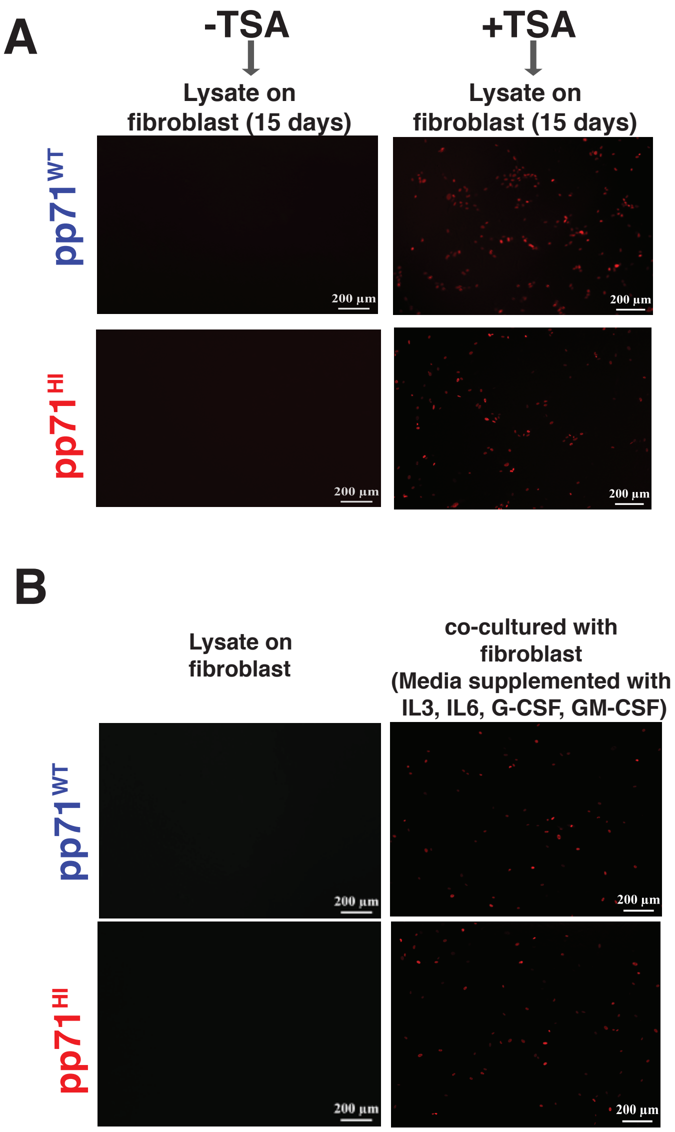


**Figure S7**: ***Reactivation of CMV in NTera2 and CD14+ cells.*** (A) Fluorescent images of HFFs treated with lysate (for 10 days) from Ntera2 cells infected with $pp71^{WT}$ or $pp71^{HI}$ virus for 4 days followed by treatment with TSA (+/-) for 24h. (B) Fluorescent images of HFFs treated with lysates (for 15 days) from $pp71^{WT}$ or $pp71^{HI}$ virus-infected CD14+ cells at 10dpi (left). At 10dpi, CD14+ cells infected with $pp71^{WT}$ or $pp71^{HI}$ virus were co-cultured with HFFs in the presence of reactivation media (media supplemented with IL3, IL6, G-CSF, GM-CSF), and imaged at 15 days post co-culturing (right).

**SUPPLEMENTARY TABLES**

Table S1: *List of probes and primers for qRT-PCR.*

| Primer | Sequence |
|---|---|
| pNL4-3 reference RT R | TGTGTGCCCGTCTGTTGTGT |
| pNL4-3 reference RT F | GAGTCCTGCGTCGAGAGATCTC |
| pNL4-3 reference probe | FAM/CAGTGGCGCCCGAACAGGGA/TAMSp/ |
| HCMV UL83 RT F | TCTTCCTGGAGGTACAAGCCA |
| HCMV UL83 RT R | CAGCCACGGGATCGTACTG |
| HCMV probe | HEX/ACGCGAGACCGTGGAACTGCG/ [black hole quencher-1] |
| UL54 F | CCCTCGGCTTCTCACAACAAT |
| UL54 R | CGAGTTAGTCTTGGCCATGCAT |
| β-actin F | CATTGCCGACGGATGCA |
| β-actin R | GCCGATCCACACGGAGTACT |
